# Functional localization of the frontal eye fields in the common marmoset using microstimulation

**DOI:** 10.1101/715359

**Authors:** Janahan Selvanayagam, Kevin D. Johnston, David J. Schaeffer, Lauren K. Hayrynen, Stefan Everling

## Abstract

The frontal eye field (FEF) is a critical region for the deployment of overt and covert spatial attention. While investigations in the macaque continue to provide insight into the neural underpinnings of the FEF, due to its location within a sulcus the macaque FEF is virtually inaccessible to electrophysiological techniques such as high-density and laminar recordings. With a largely lissencephalic cortex, the common marmoset (*Callithrix jacchus*) is a promising alternative primate model for studying FEF microcircuitry. Putative homologies have been established with the macaque FEF on the basis of cytoarchitecture and connectivity, however physiological investigation in awake, behaving marmosets is necessary to physiologically locate this area. Here we addressed this gap using intracortical microstimulation in a broad range of frontal cortical areas in marmosets. We implanted marmosets with 96-channel Utah arrays and applied microstimulation trains while they freely viewed video clips. We evoked short-latency fixed vector saccades at low currents (<50 μA) in areas 45, 8aV, 8C and 6DR. We observed a topography of saccade direction and amplitude consistent with findings in macaques and humans; we observed small saccades in ventrolateral FEF and large saccades combined with contralateral neck and shoulder movements encoded in dorsomedial FEF. Our data provide compelling evidence supporting homology between marmoset and macaque FEF and suggest the marmoset is a useful primate model for investigating FEF microcircuitry and its contributions to oculomotor and cognitive functions.

**Significance Statement:** The frontal eye field (FEF) is a critical cortical region for overt and covert spatial attention. The microcircuitry of this area remains poorly understood, as in the macaque, the most commonly used model, it is embedded within a sulcus and is inaccessible to modern electrophysiological and optical imaging techniques. The common marmoset is a promising alternative primate model due to its lissencephalic cortex and potential for genetic manipulation. However, evidence for homologous cortical areas in this model remains limited and unclear. Here we applied microstimulation in frontal cortical areas in marmosets to physiologically identify the FEF. Our results provide compelling evidence for a frontal eye field in the marmoset, and suggest that the marmoset is a useful model for FEF microcircuitry.

## Introduction

Described originally by Ferrier (1875) as a cortical area in macaque monkeys where electrical stimulation elicited contralateral eye and head movements, the frontal eye fields (FEF) in macaques and humans are now increasingly regarded as not only a motor area for saccades and head movements, but also as a critical region for the deployment of overt and covert spatial attention (Awh et al., 2006). Over the past 40 years, most of our knowledge regarding the neural processes in the FEF has come from experiments in awake behaving macaque monkeys. In these Old-World primates, FEF is defined as an area within the rostral bank and fundus of the arcuate sulcus from which electrical microstimulation evokes saccades at low currents (<50 μA) (Bruce et al., 1985). Stimulation, recording, and pharmacological manipulation studies in trained macaque monkeys have and continue to provide critical insights into the neural processes in FEF that underlie saccade control and visual attention. However, the local FEF microcircuitry remains poorly understood as, due to its location within a sulcus, macaque FEF is virtually inaccessible to intralaminar recordings and manipulations.

The New-World common marmoset (*Callithrix jacchu*s) is a promising alternative primate model for studying FEF microcircuitry. These small primates have a largely lissencephalic cortex and can be trained to perform saccadic eye movement tasks head-restrained (Mitchell et al., 2014; Johnston et al., 2018, 2019). A first step towards such experiments is the physiological identification of the FEF in marmosets. Existing evidence for the location of this area in this species, however, remains limited and unclear. An early marmoset study by Mott and colleagues (1910) reported that both eye and combined eye and head movements could be evoked by electrical stimulation at several frontal cortical sites. Subsequently, Blum and colleagues (1982) confirmed and extended these earlier results. They observed movements including ipsilateral and contralateral saccades, eye movements in all directions, and slow drifting movements. It seems that these eye movements were evoked in areas 6DC, 6DR, 8aD, and 46 with no clear topography of direction or amplitude. Interpretation of these earlier studies is difficult, however, as the anesthetized preparations used most likely influenced the properties of the eye movements evoked (Robinson and Fuchs, 1969).

More recently, anatomical evidence has suggested that marmoset FEF lies within areas 45 and 8aV (Reser et al., 2013). Both areas have widespread connections with extrastriate visual areas, and areas labelled FEF and FV by Collins et al (2005), which may correspond to areas 45 and 8aV, contain clusters of neurons projecting to the SC, an area critical for the initiation of saccadic and orienting movements. Area 8aV in marmosets also contains large layer V pyramidal neurons, a cytoarchitectonic characteristic of macaque FEF (Stanton et al., 1989). Consistent with this notion, fMRI studies in marmosets have reported BOLD activation in areas 45 and 8aV in response to visual stimuli (Hung et al., 2015), though a resting-state fMRI functional connectivity study found the strongest SC connectivity in area 8aD, at the border of area 6DR (Ghahremani et al., 2017). The authors proposed that this region either corresponded to the marmoset FEF or that it may encode large amplitude saccades, while area 8aV may encode small amplitude saccades.

Here, we set out to physiologically identify the marmoset FEF using the classical approach of intracortical electrical microstimulation (ICMS). We applied microstimulation trains via chronically implanted 96-channel electrode arrays placed to target a broad range of frontal cortical areas in three awake marmosets. Our findings revealed a topography of contralateral saccade amplitude in marmoset frontal cortex similar to that observed in macaques (Bruce et al., 1985; Schall, 1997) and humans (Foerster, 1926), with small saccades being encoded in area 45 and lateral parts of area 8aV, and larger saccades combined with contralateral neck and shoulder movements encoded in the medial posterior portion of area 8aV, area 8C, and area 6DR.

## Methods

### Subjects

We obtained data from 3 adult common marmosets (*Callithrix jacchus*; M1 male, 17 months; M2 female 20 months; M3 male 23 months). All experimental procedures conducted were in accordance with the Canadian Council of Animal Care policy on the care and use of laboratory animals and a protocol approved by the Animal Care Committee of the University of Western Ontario Council on Animal Care. The animals were under the close supervision of university veterinarians.

Prior to the commencement of microstimulation experiments, each animal was acclimated to restraint in a custom primate chair (Johnston et al., 2018). Animals then underwent an aseptic surgical procedure under general anesthesia in which 96 channel Utah arrays (4mm × 4mm; 1mm electrode length; 400μm pitch; iridium oxide tips) were implanted in left frontal cortex. During this surgery, a microdrill was used to initially open 4mm burr holes in the skull and were enlarged as necessary using a rongeur. Arrays were manually inserted; wires and connectors were fixed to the skull using dental adhesive (Bisco All-Bond, Bisco Dental Products, Richmond, BC, Canada). Once implanted, the array site was covered with silicone adhesive to seal the burr hole (Kwik Sil, World Precision Instruments, Sarasota, FLA, USA). A screw-hole was drilled into the skull on the opposite side to the location of the implanted array to place the ground screw. The ground wire of the array was then tightly wound around the base of the screw to ensure good electrical connection. A combination recording chamber/head holder (Johnston et al., 2018) was placed around the array and connectors and fixed in place using further layers of dental adhesive. Finally, a removable protective cap was placed on the chamber.

### Localizing the array

To precisely determine array locations, high-resolution T2-weighted structural magnetic resonance images (MRI; obtained pre-surgery) were co-registered with computerized tomography (CT) scans (obtained post-surgery). The MRI images provided each marmoset’s brain geometry with reference to the location of the skull, while the CT images allowed for localization of the skull and the array boundaries. By co-registering the skulls across the two modalities, the precise array-to-brain location was determined for each animal.

Pre-surgical MRIs were acquired using an 9.4 T 31 cm horizontal bore magnet (Varian/Agilent, Yarnton, UK) and Bruker BioSpec Avance III console with the software package Paravision-6 (Bruker BioSpin Corp, Billerica, MA) and a custom-built high performance 15-cm-diameter gradient coil with 400-mT/m maximum gradient strength (xMR, London, CAN; Peterson et al., 2018). A geometrically optimized 8-channel phased array receive coil was designed in-house, for SNR improvement and to allow for acceleration of the echo planar imaging of marmoset cohorts (Gilbert et al., 2019). Preamplifiers were located behind the animal and the receive coil was placed inside a quadrature birdcage coil (12-cm inner diameter) used for transmission. Prior to each imaging session, anesthesia was induced with ketamine hydrochloride at 20 mg/kg. During scanning, marmosets were anesthetized with isoflurane and maintained at a level of 2% throughout the scan by means of inhalation. Oxygen flow rate was kept between 1.75 and 2.25 l/min throughout the scan. Respiration, SpO2, and heart rate were continuously monitored and were observed to be within the normal range throughout the scans. Body temperature was also measured and recorded throughout, maintained using warm water circulating blankets, thermal insulation, and warmed air. All animals were head-fixed in stereotactic position using a custom-built MRI bed with ear bars, eye bars, and a palate bar housed within the anesthesia mask (Gilbert et al., 2019). All imaging was performed at the Centre for Functional and Metabolic Mapping at the University of Western Ontario. T2-weighted structural scans were acquired for each animal with the following parameters: TR = 5500 ms, TE = 53 ms, field of view = 51.2 × 51.2 mm, matrix size = 384 × 384, voxel size = 0.133 × 0.133 × 0.5 mm, slices = 42, bandwidth = 50 kHz, GRAPPA acceleration factor: 2.

CT scans were obtained on a micro-CT scanner (eXplore Locus Ultra, GR Healthcare Biosciences, London, ON) after array implantation. Prior to the scan, marmosets were anesthetized with 15mg/kg Ketamine mixed with 0.025mg/kg Medetomidine. X-ray tube potential of 120 kV and tube current of 20 mA were used for the scan, with the data acquired at 0.5º angular increment over 360º, resulting in 1000 views. The resulting CT images were then reconstructed into 3D with isotropic voxel size of 0.154 mm. Heart rate and SpO2 were monitored throughout the session. At the end of the scan, the injectable anesthetic was reversed with an IM injection of 0.025mg/kg Ceptor.

The raw MRI and CT images were converted to NifTI format using dcm2niix (Li et al., 2016) and the MRIs were reoriented from the sphinx position using FSL software (Smith et al., 2004). Then, using FSL (FSLeyes nudge function), each animal’s CT image was manually aligned to their MRI image based on the skull location – this allowed for co-localization of the array and brain surface. The array position from the CT image was determined by a hyper-intensity concomitant with the metallic contacts contained within the array; this hyper-intensity stood out against the lower intensities of the skull and surrounding tissues. A region of interest (ROI) was manually drawn within the array location for each animal to be displayed on the NIH marmoset brain atlas surface (Liu et al., 2018) for ease of viewing. The NIH marmoset brain atlas is an ultra-high resolution *ex vivo* MRI image dataset that contains the locations of cytoarchitectonic boundaries (Liu et al., 2018). As such, to determine the array location with reference to the cytoarchitectonic boundaries, we non-linearly registered the NIH template brain to each marmoset’s T2-weighted image using Advanced Normalization Tools (ANTs; Avants et al., 2011) software. The resultant transformation matrices were then applied to the cytoarchitectonic boundary image included with the NIH template brain atlas. The olfactory bulb was manually removed from the marmoset T2-weighted image of each animal prior to registration, as it was not included in the template image. As a result of the transformations, the template brain surface, the cytoarchitectonic boundaries, and the array location (ROI described above) could be rendered on each animals’ individual native-space brain surface.

### Data collection

Following recovery, we verified that electrode contacts were within the cortex by monitoring extracellular neural activity using the Open Ephys acquisition board (http://www.open-ephys.org) and digital headstages (Intan Technologies, Los Angeles, CA, USA). Upon observing single or multiunit activity at multiple sites in the array, we commenced microstimulation experiments.

Animals were head restrained in a custom primate chair (Johnston et al., 2018) mounted on a table in a sound attenuating chamber (Crist Instruments Co., Hagerstown, MD, USA). A spout was placed at the monkey’s mouth to deliver a viscous preferred reward of acacia gum. This was delivered via infusion pump (Model NE-510, New Era Pump Systems, Inc., Farmingdale, New York, USA). In each session, eye position was calibrated by rewarding 300 to 600ms fixations on a marmoset face presented at one of five locations on the display monitor using the CORTEX real-time operating system (NIMH, Bethesda, MD, USA). Faces were presented at the display centre, at 6 degrees to the right and left of centre, and at 6 degrees directly above and below centre. All stimuli were presented on a CRT monitor (ViewSonic Optiquest Q115, 76 Hz non-interlaced, 1600 × 1280 resolution).

Monkeys freely viewed short repeating video clips to sustain their alertness while we applied manually triggered microstimulation trains. Monkeys were intermittently rewarded at random time intervals to maintain their interest. Microstimulation trains were delivered using the Intan RHS2000 Stimulation/Recording Controller system and digital stimulation/recording headstages (Intan Technologies, Los Angeles, CA, USA). Stimulation trains consisted of 0.2-0.3ms biphasic current pulses delivered at 300 Hz for a duration of 100-400ms, at current amplitudes varying between 5 and 300 μA. At sites where skeletomotor or saccadic responses were evoked, we carried out a current series to determine thresholds. The threshold was defined as the minimum current at which a given response was evoked on 50% of stimulation trials. Skeletomotor responses were observed manually by researchers. Eye position was digitally recorded at 1 kHz via video tracking of the left pupil (EyeLink 1000, SR Research, Ottawa, ON, Canada).

### Data analysis

Analysis was performed with custom python code. Eye velocity (visual deg/s) was obtained by smoothing and numerical differentiation. Saccades were defined as horizontal or vertical eye velocity exceeding 30 deg/s. Blinks were defined as the radial eye velocity exceeding 1500 deg/s.

As we did not require marmosets to fixate during stimulation, saccades following stimulation could be spontaneous. A bootstrap analysis was used to quantitatively determine if saccades were more probable following stimulation than at any other time during a session. In a single session, 60-80 trains were delivered at a single site holding stimulation parameters constant over a 2-minute period. Stimulation onset times were shuffled (time points were randomly sampled without replacement with millisecond resolution over the duration of the session) and the probability of a saccade occurring in a 200ms window following the selected timepoints was computed. This was repeated 1000 times for each session to obtain a distribution of probabilities of saccade occurrence. The percentile rank of the probability of stimulation evoking a saccade with respect to this distribution was computed; the 95th percentile marked the 5% significance criterion indicating a session where stimulation significantly increased the probability of saccade occurrence.

## Results

### Evoked skeletomotor and oculomotor responses

Array locations were confirmed using CT scans obtained after the surgery, which were co-registered with MR scans obtained before the surgery (see Fig. 1a). Microstimulation was conducted at 288 sites across 3 marmosets. We observed a range of skeletomotor and oculomotor responses across the frontal cortex (Fig. 1b, c).

**Fig 1.**
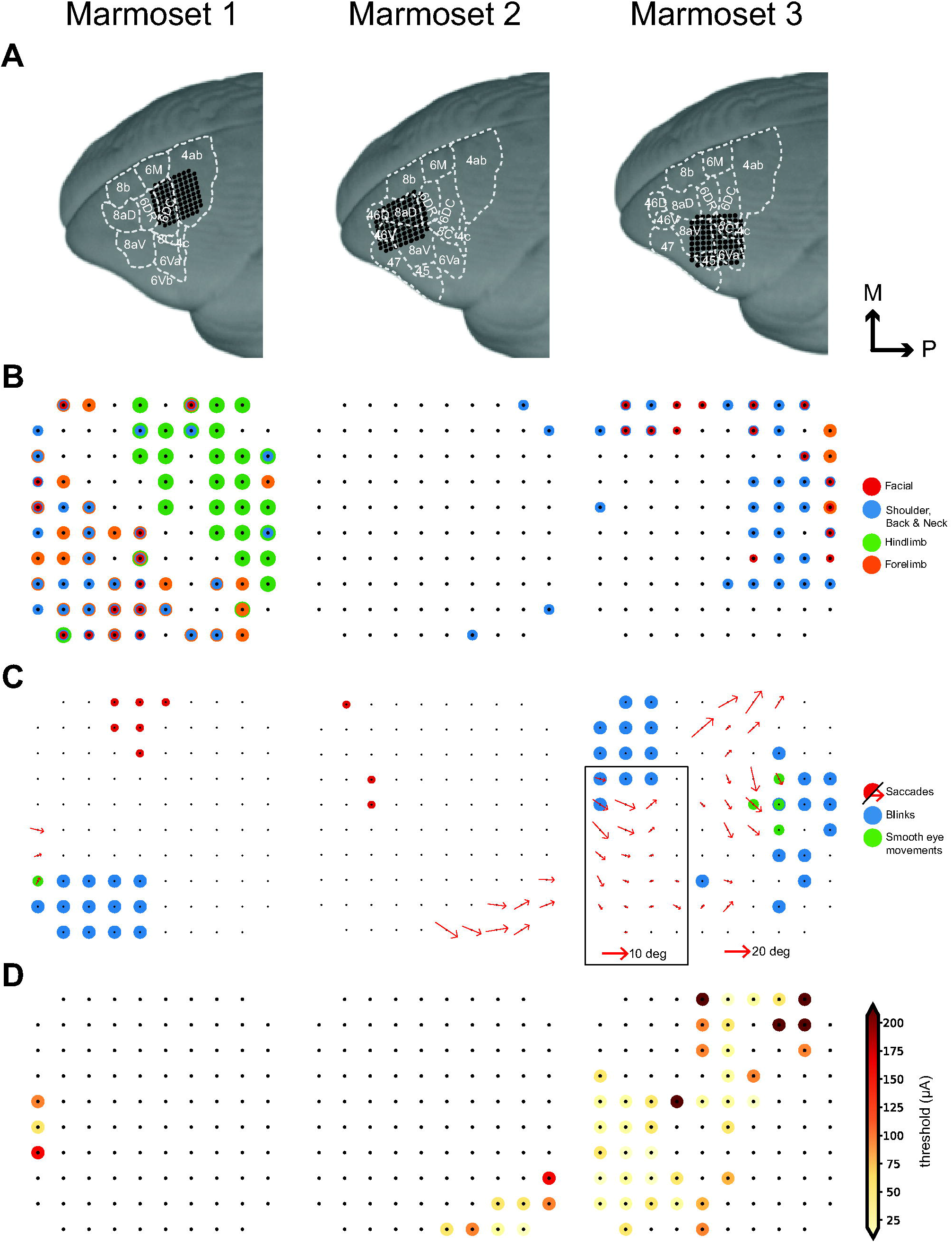
Evoked motor responses. (A) Array locations in each marmoset reconstructed using MR and CT images (see <u>Localizing the array</u>). (B) Pattern of evoked skeletomotor responses in each marmoset. (C) Pattern of evoked oculomotor responses in each marmoset. At sites where fixed vector saccades were observed, mean saccade vector is plotted. Mean saccade vectors were computed at the minimum current where saccades are evoked at least 75% of the time. Inset shows small saccade vectors at 2x scale for Marmoset 3. (D) Thresholds for saccades at sites where saccades were evoked at currents <= 300μA.

At the most posterior sites, we observed primarily single joint movements with a gross medio-lateral topography. We observed hindlimb movements (leg, foot, toes) most medially, followed by forelimb (arm, hand, finger) and facial movements (eyelid, ear, nose, jaw) most laterally - an organization characteristic of primary motor cortex (area 4) (Burish et al., 2008; Wakabayashi et al., 2018). Anterior to this, we observed overlapping representation of forelimb, facial, shoulder, and neck musculature with no obvious organization, similar to that observed in the marmoset premotor cortex (area 6) (Burish et al., 2008; c.f. Wakabayashi et al., 2018).

We elicited saccades at 61 sites across 3 marmosets (see Fig. 1c). At 6 sites on the border of area 6DC and 6M, we observed goal directed saccades characteristic of the supplementary eye fields (SEF), albeit at long latencies (70-110ms) and high currents (200 μA) (see Fig. 3a). At 3 sites in area 46D and the anterior portion of area 8aD, we elicited saccades with no clear pattern at long latencies (75-90ms) and high currents (300 μA) (see Fig. 3b). Saccades evoked from these sites were mostly directed to the hemifield contralateral to the stimulated site, though some saccades directed to the ipsilateral hemifield were observed.

We elicited fixed vector saccades at 52 sites across areas 6DR, 8C, 8aV and 45. Mean saccade vectors are plotted in Fig. 1c. Representative saccade traces are plotted in Fig. 2. In areas 6DR, 8C and the medial portion of 8aV, we observed larger saccades often coupled with shoulder, neck, and ear movements with the most common response being a shoulder rotation that resembled orienting towards contralateral side. In area 45 and the lateral portion of area 8aV, we observed smaller saccades with no visible skeletomotor responses. Smooth eye movements could be elicited at 5 sites in areas 6DR and 8C.

**Fig 2.**
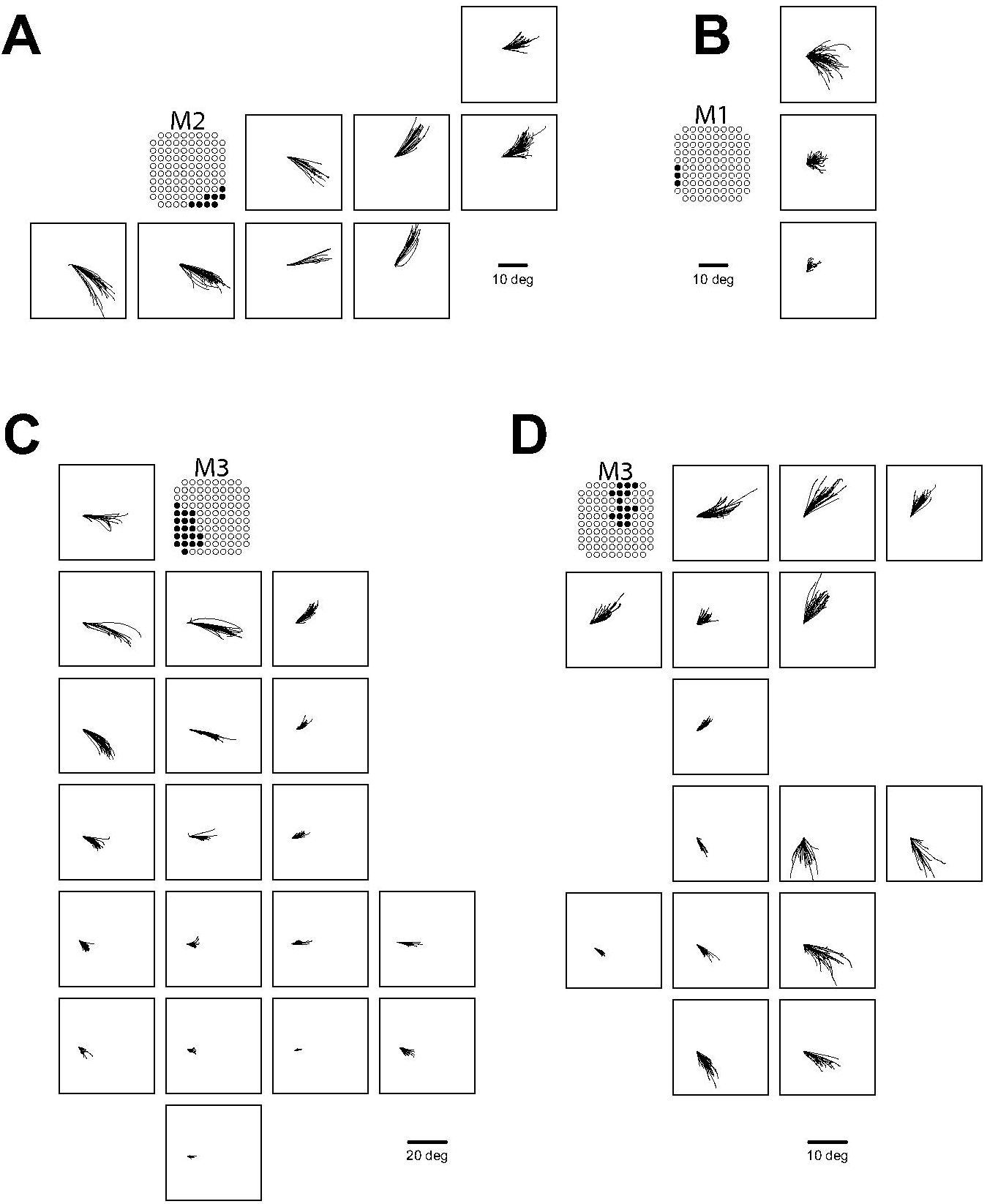
Saccades evoked in FEF sites. Representative traces for fixed vector saccades in (A) Marmoset 2 (A), Marmoset 1 (B) and Marmoset 3 (C, D).

**Fig 3.**
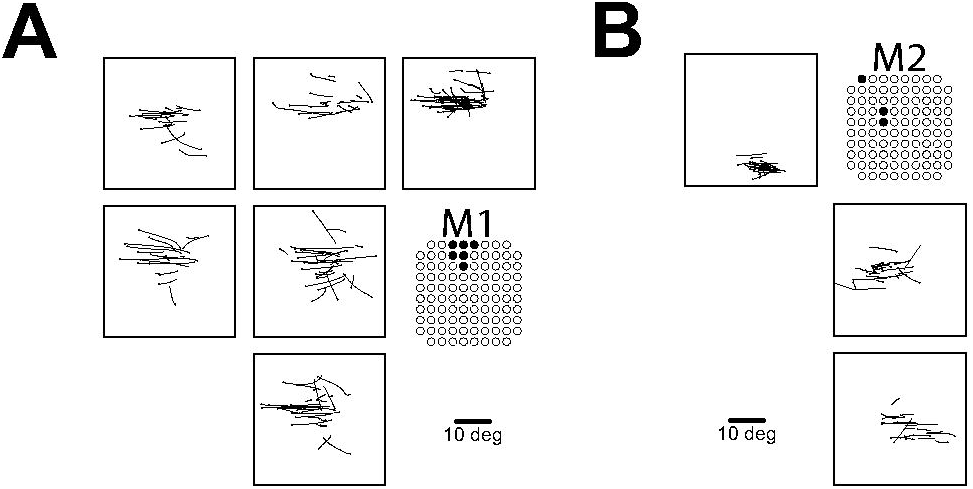
Saccades evoked in non-FEF sites. Representative traces for goal-directed saccades from dorsomedial sites in Marmoset 1 (A) and saccades from rostral sites in Marmoset 2 (B). Open circles indicate eye position at saccade onset.

### Saccade thresholds and latencies

At sites where we observed fixed vector saccades, we conducted current series to determine thresholds and characterize any current-related changes in saccade metrics. Current series from five representative sites are shown in Fig. 4a-e. Thresholds were defined as the minimum current at which saccades could be evoked 50% of the time (see Fig. 4g). Thresholds ranged from 12-300 μA. Saccades were evoked at low thresholds (<50 μA) at 35 of the 52 sites from which we were able to evoke fixed vector saccades (see Fig. 1d). Saccade metrics were computed at the minimum current at which saccades could be evoked 75% of the time.

**Fig 4.**
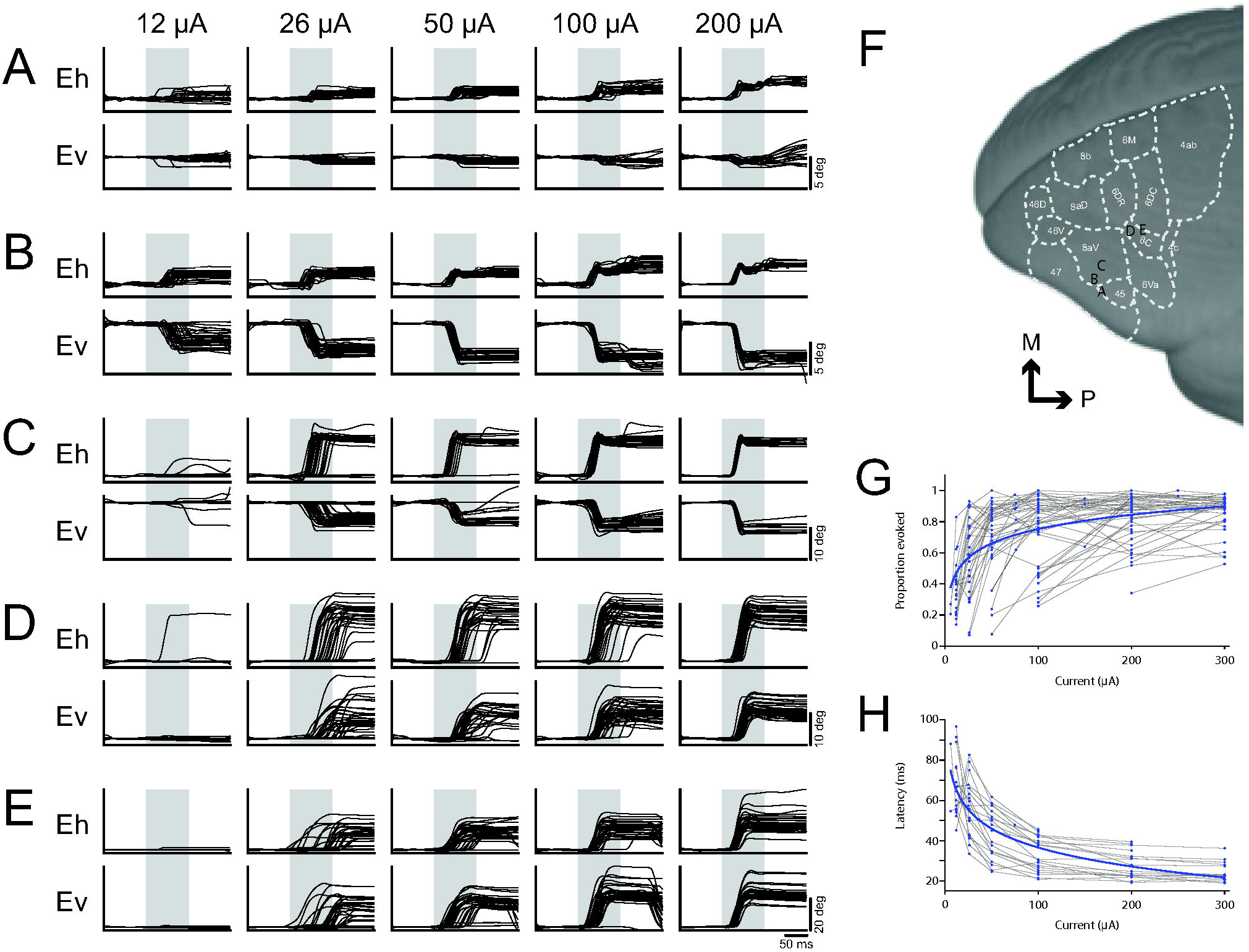
Current series at representative saccade sites. Current series at a representative small (A-C) and large (D-E) saccade sites. Grey bars indicate stimulation train duration. Location of array sites for series in (A-E) show in (F). (G) Effect of current on proportion of saccades evoked at all FEF sites in Marmoset 3. (H) Effect of current on saccade latency at low threshold (<50 μA) sites in Marmoset 3.

Each site had a stereotypical saccade latency, though we found no systematic variation in saccade latency with respect to site coordinates nor any other saccade metrics. Saccade latencies ranged from 25-85ms, with the majority falling in the range between 40-60ms (see Fig. 4h). Saccade latencies were generally longer and more variable near the current threshold for a given site. When using high currents well above threshold (200-300 μA), uniformly short saccade latencies were observed (15-45ms).

### Topography of evoked saccades

Evoked saccades were directed contralateral to the stimulated hemisphere and mostly fixed vector (see Fig 1c, Fig 2, Fig 4a-e), exhibiting relatively consistent directions and amplitudes independent of the initial eye position. Although we did not systematically vary initial eye positions, the fact that marmosets were allowed to freely direct their gaze across video clips on the display monitor during experimental sessions ensured a wide range of initial eye positions at the time of microstimulation onset. Most initial eye positions fell within a 13 degree range similar to observations elsewhere in marmosets (Mitchell et al., 2014) and other New World monkeys (Heiney and Blazquez, 2011). 90% of initial eye positions fell within the following ranges for each marmoset: Marmoset 1: −13.6 to 12.4 abscissa, −10.7 to 11.4 ordinate; Marmoset 2: −12.7 to 15.7 abscissa, −11.7 to 9.6 ordinate; Marmoset 3: −12.9 to 12.7 abscissa, −18.5 to 14.3 ordinate. Amplitude decreased progressively from medial (large saccades; >20 visual degrees) to lateral (small saccades; <2 visual degrees) sites. Direction varied systematically from upper visual field at posterior medial sites to lower visual field at anterior lateral sites.

### Staircase saccades

At a subset of sites from which saccades were evoked, we additionally observed staircases of multiple saccades. To investigate this further, we applied stimulation trains of increasing duration at these sites and found that the number of saccades increased as a function of train duration at the majority of these sites (12/15). A representative site is depicted in Fig. 5. Staircases consisted of 2-5 consecutive saccades with consistent amplitudes and directions, in many cases ultimately driving the eye to the extent of its oculomotor range. At a given site, consecutive saccades occurred at fixed intervals. The intersaccadic interval ranged from 70-120 ms across sites and we observed no systematic variation in intersaccadic interval with respect to site coordinates nor any other saccade metrics.

**Fig 5.**
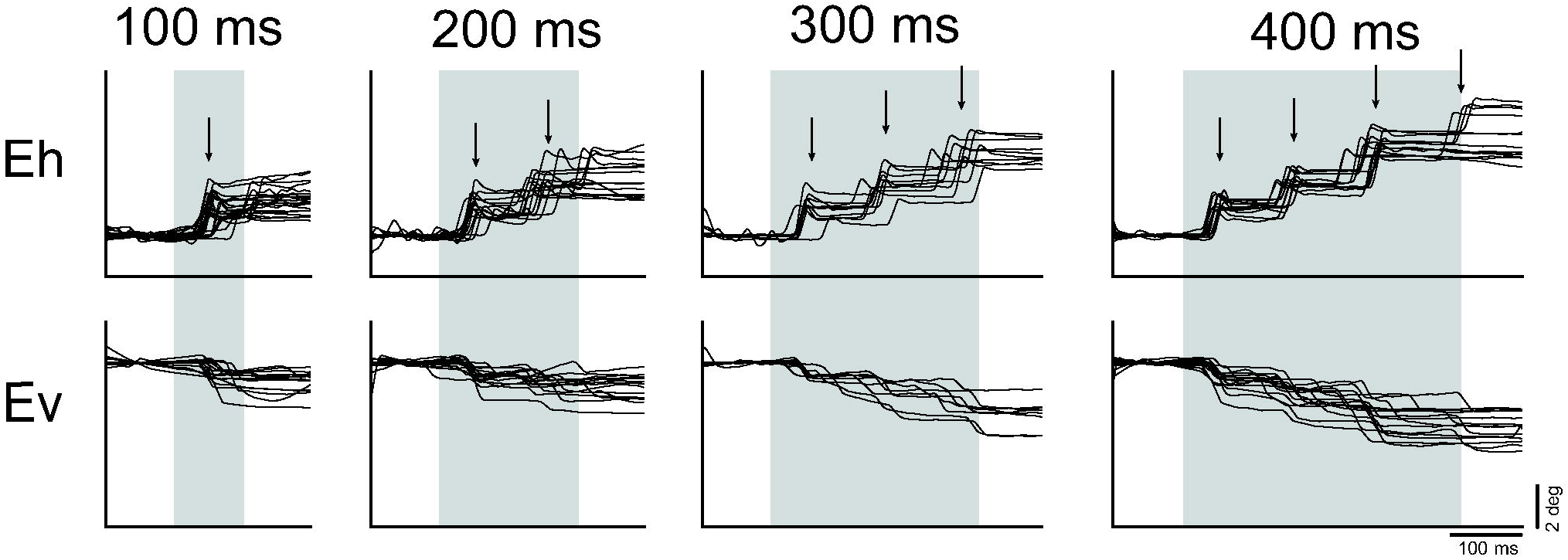
Current series at a representative site with staircase saccades. Arrows indicate median saccade onset latency. Grey bars indicate stimulation train duration.

### Smooth eye movements

Posterior to where we evoked saccades, in areas 6DR and 8C (see Fig. 1c), we were able to elicit smooth eye movements. These eye movements often followed a saccade and continued until stimulation ended at which point, they stopped abruptly (see Fig. 6a for a representative site). While the direction of these movements was consistent at a site, the velocity increased as a function of stimulation current intensity, consistent with what is observed in the smooth pursuit region of the FEF in macaques (see Fig. 6b for a current series at a representative site) (Gottlieb et al., 1993).

**Fig 6.**
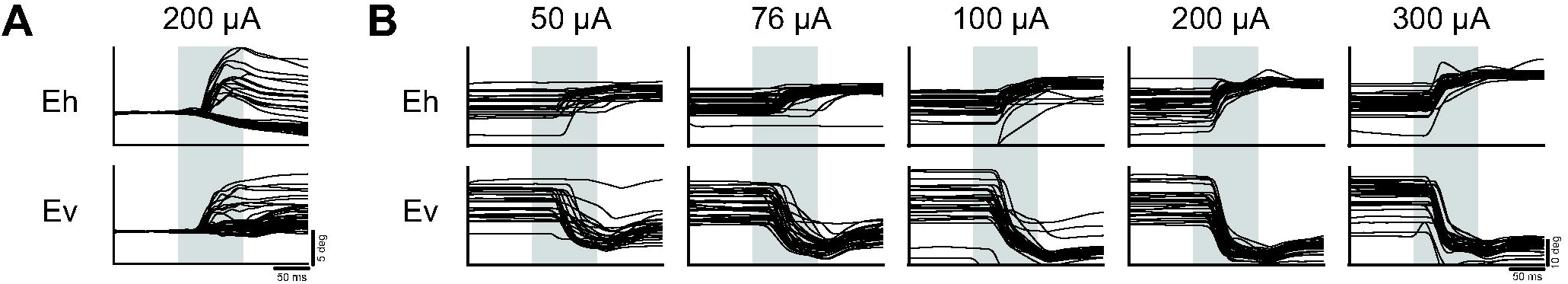
Evoked smooth eye movements. (A) Smooth eye movement site at 200 μA from Marmoset 1. (B) Current series from a smooth eye movement site in Marmoset 3. Grey bars indicate stimulation train duration.

### Effects of initial gaze position

While evoked saccades were mostly fixed vector, an effect of initial gaze position was observed at some sites. At those sites, saccades tended to be of greater amplitude if the gaze position at the time of stimulus onset was within the hemifield ipsilateral to the stimulated hemisphere. Further, the probability of evoking a saccade was lower if the initial eye position was within the hemifield contralateral to the stimulated hemisphere.

We quantified the magnitude of the effect of initial eye position at each site by computing the linear regression of the difference in final eye position as a function of the initial eye position separately for horizontal (*K*_*h*_) and vertical (*K*_*v*_) components of evoked saccades at these sites (Russo and Bruce, 1993). Correlation coefficients of 0 would be expected for sites at which the saccade vector did not change with varying initial eye positions (i.e. strictly fixed-vector saccades), whereas coefficients of −1 would be expected for sites at which evoked saccades terminated at the same eye position irrespective of initial eye position (i.e. goal-directed saccades). An example of this is shown for representative sites from FEF (see Fig. 6a, b) and SEF (see Fig. 6c).

Sites in FEF were mostly fixed vector, however, as observed by Russo and Bruce (1993), the effect of initial eye position increases in magnitude with the mean amplitude of saccades evoked at that site (see Fig. 6d). This corresponds with the eye position terminating at the edge of the orbit for very large saccades. In contrast, in SEF sites, mostly convergent saccades were observed with correlation coefficients close to −1 and saccades converging on locations well within the oculomotor range of the animal.

## Discussion

The common marmoset is a promising model for investigating the microcircuitry of the FEF (Mitchell and Leopold, 2015). The location of the FEF in marmosets, however, remains controversial. To address this, we systematically applied intracortical microstimulation (ICMS) to marmoset frontal cortex through chronically implanted electrode arrays to investigate the oculomotor and skeletomotor responses evoked in this region (see Fig. 7 for a schematic summary). We observed patterns of skeletomotor responses consistent with previous ICMS investigations of marmoset motor and premotor cortex (Burish et al., 2008; Wakabayashi et al., 2018). Anterior to these motor areas, we observed a suite of oculomotor responses across frontal cortex which we propose correspond to three cortical eye fields. ICMS in area 45 and in the lateral part of area 8aD evoked small contraversive saccades at very low currents, consistent with the properties of the ventrolateral FEF (vFEF) in macaques (Bruce et al. 1985). In areas 6DR, 6DC, 8C, and medial 8aV, ICMS evoked larger saccades that were often associated with shoulder, neck and ear movements. This is consistent with ICMS experiments in dorsomedial macaque FEF (dFEF) (Elsley et al., 2007; Corneil et al., 2010). We also observed goal-oriented saccades characteristic of the supplementary eye field (SEF) at dorsomedial sites. In prefrontal areas 46 and anterior 8aD, ICMS elicited saccades with no consistent organization of direction or amplitude. These findings are consistent with the organization of FEF and SEF in macaques (Robinson and Fuchs, 1969; Bruce et al., 1985; Schlag and Schlag-Rey, 1987; Gottlieb et al., 1993; Russo and Bruce, 1993; Knight and Fuchs, 2007).

**Fig 7.**
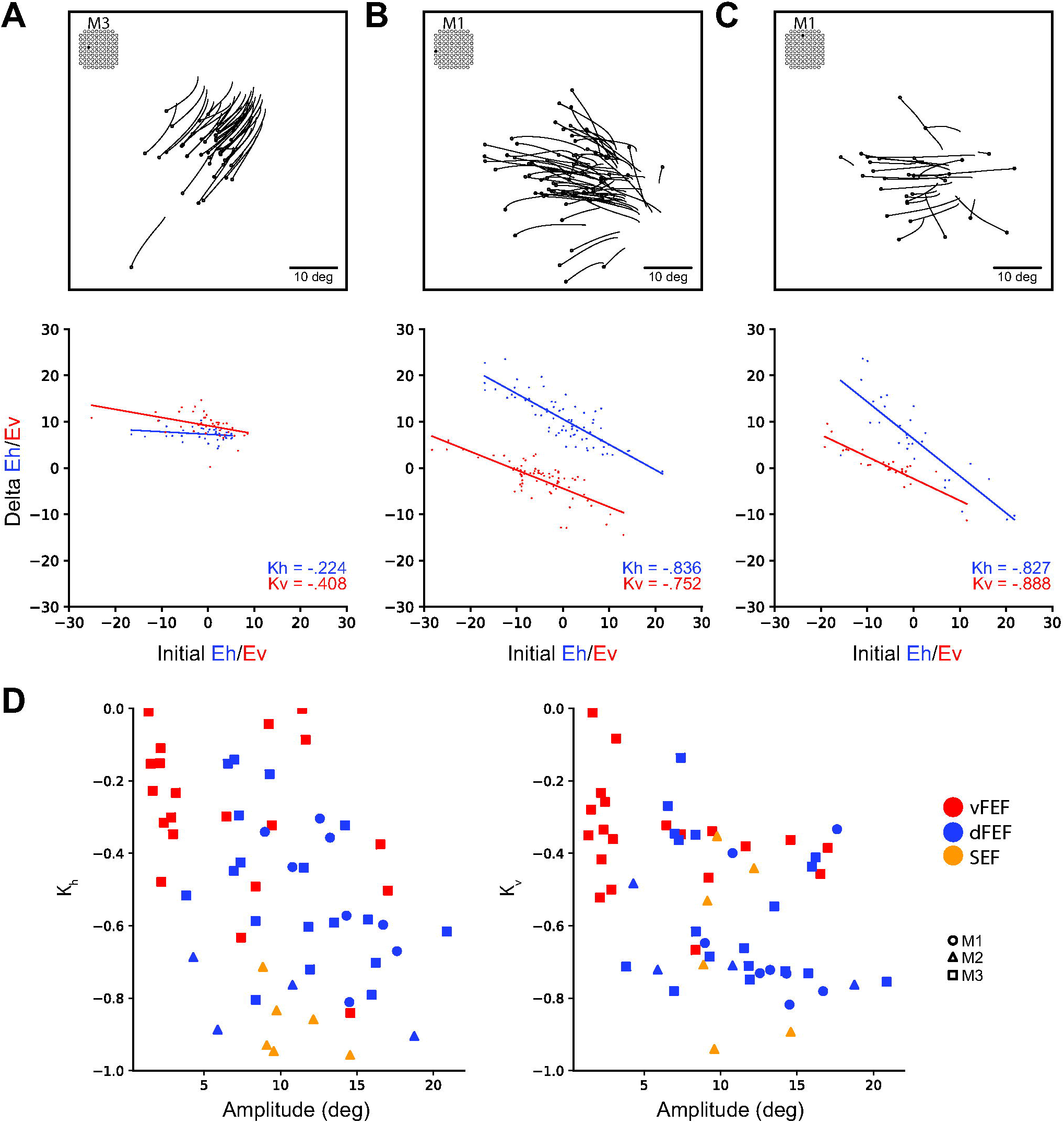
Effect of initial eye position. Saccade traces (above) and effect of initial position on delta (below) for representative sites from vFEF (A), dFEF (B) and SEF (C). Open circles indicate eye position at saccade onset. (C) Across all sites, the relationship between *K*_*h*_ and *K*_*v*_ values (correlation coefficients from effect of initial eye position analysis) and amplitude. More negative values indicate a greater effect of initial eye position.

**Fig 8.**
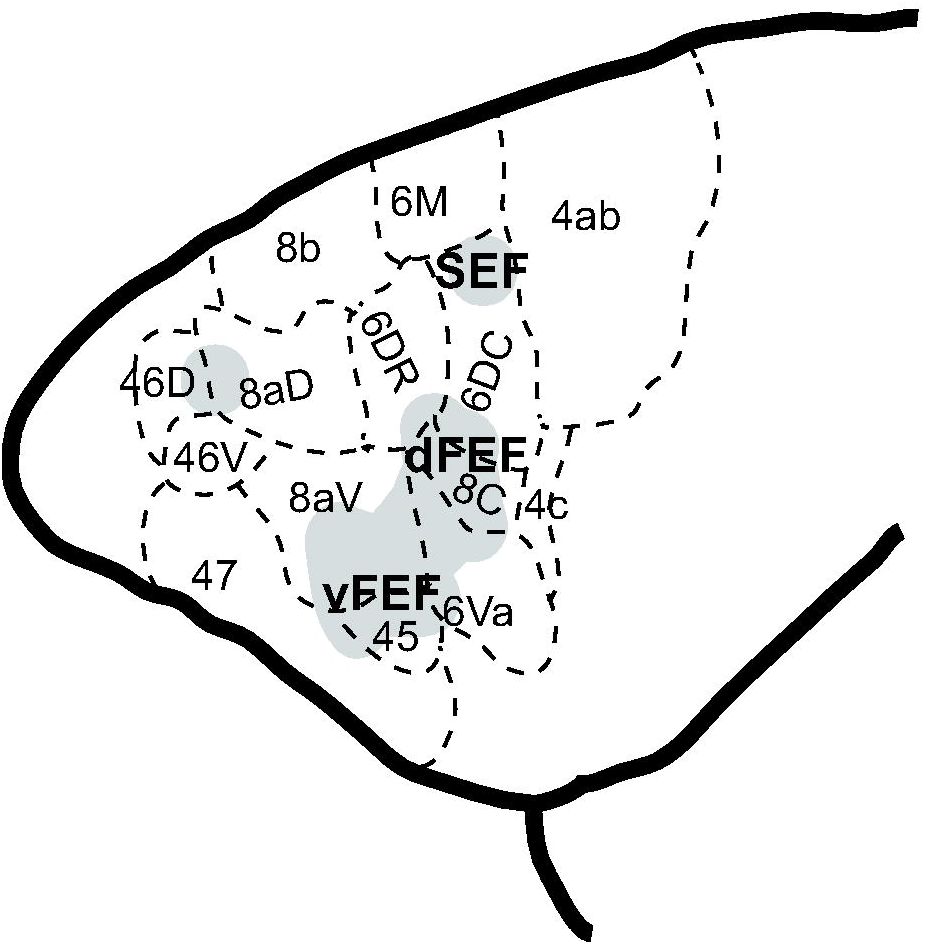
Schematic representation of cortical eye fields in marmoset frontal cortex.

A characteristic feature of the FEF observed in macaque ICMS experiments is the ability to evoke short latency fixed vector saccades at low currents. While the threshold to evoke saccades can be as high as 2 mA in frontal cortex (Robinson and Fuchs, 1969), FEF is defined in macaque as the restricted region in which thresholds are below 50 μA (Bruce et al., 1985). Here, we observed a large number of sites with thresholds below 50 μA, with a lower bound of 12 μA, similar to the 10 μA observed in macaque (Bruce et al., 1985). This is despite the limitations of fixed-length chronic electrode arrays which did not allow us optimally target layer V output neurons and in contrast to previous reports of higher thresholds in marmoset motor cortex compared to macaques (Burish et al., 2008). However, saccade latencies were slightly longer than those observed in macaques. We found a range of 25-85ms as compared to 20-60ms observed by Bruce and colleagues (1985) at near threshold currents, and 15-45ms as compared to 15-25ms by Robinson and Fuchs (1969) at higher currents. It has been proposed that longer latency saccades are evoked through an indirect route (e.g., superior colliculus), whereas shorter latency saccades are evoked by recruiting neurons that project directly to the brain stem (Bruce et al., 1985). Investigations employing single unit recordings in the marmoset FEF and studies investigating the connectivity of marmoset FEF and brain stem oculomotor nuclei should provide insight into these differences.

In macaque FEF, saccades evoked by ICMS are fixed-vector with little variability in amplitude and direction (Robinson and Fuchs, 1969; Bruce et al., 1985). While saccades evoked here were predominantly fixed vector, some effects of initial gaze position were observed in which saccades were larger when the initial gaze position was in the hemifield ipsilateral to the site of stimulation. Similar observations have been made in macaque FEF (Robinson and Fuchs, 1969; Russo and Bruce, 1993) in which the magnitude of this effect is greater for larger saccades. However, this effect is greater here than previously observed with macaques. This may be a result of the eye being driven to the edge of the oculomotor range. In marmosets, this is limited to approximately 12 degrees as compared to 30 degrees in the macaque (Tomlinson and Bahra, 1986; Heiney and Blazquez, 2011; Mitchell et al., 2014). Head-restraint also prevents marmosets from using head movements to shift gaze, which they depend on to a greater extent than larger primates (Mitchell et al., 2014). Investigations in head unrestrained marmosets would clarify these differences.

Previous studies of macaque FEF have revealed a topographic representation of saccade amplitude and direction. Bruce and colleagues (1985) demonstrated a medio-lateral gradient in which large saccades were evoked medially and small saccades laterally. We observed a similar organization of saccade amplitude in marmosets, with small saccades being elicited in areas 45 and lateral area 8aV (vFEF) and larger saccades being evoked in areas 6DR, 6C, 8C, and medial 8aV (dFEF). Bruce and colleagues (1985) observed systematic changes in saccade direction with small advances along the depth of the arcuate sulcus in macaques, though they often encountered disruptions and reversals of direction. We observed a rostro-caudal organization of saccade direction in marmosets in which direction gradually changed from lower to upper visual field, though there were occasional direction reversals. Assuming that frontal cortex in marmoset is roughly a flattened version of that in macaque, the rostro-caudal axis would correspond roughly to traversing the depth of the arcuate sulcus from lip to fundus in macaques. We additionally observed a more continuous medio-lateral organization of saccade direction, such that the upper visual field was represented medially. This organization would be difficult to observe in the macaque FEF due to its more complex morphology.

At more posterior-medial sites where larger saccades are represented (dFEF), we observed skeletomotor responses resembling an orienting response while we only observed oculomotor responses at the more anterior-lateral sites. This is in line with what Knight and Fuchs (2007) found in awake head-unrestrained macaques. Indeed, Foerster (1926) already reported two saccade-related fields in humans: (1) FEF where epileptic seizures evoked contralateral saccades and (2) a more posterior field that he termed frontal adversive field (frontales Adversivsfeld) where seizures were associated with contralateral saccades and head movements.

At posterior medial sites, at the border of area 6D and 6M, we observed goal-directed saccades characteristic of SEF (Schlag and Schlag-Rey, 1987). Contrary to observations at more anterior lateral sites, convergence of saccades could not be explained by physical limitation of the orbit. We observed saccades converging at locations well within the animal’s oculomotor range and, albeit infrequently, saccades directed to the hemifield ipsilateral to the stimulated hemisphere. These findings are similar to observations in the macaque by Schlag and Schlag-Rey (1987). However, we observed that saccade latencies were much longer at these sites (70-110ms) than those observed by Schlag and Schlag-Rey (1987) (40-60ms). Further, they observed low current thresholds, at many sites less than 20 μA, whereas we observed few saccades at currents as high as 200 μA. Taken together, these findings suggest the observed responses may be evoked due to current spread to dorsomedial regions not covered by our arrays. We propose that area 6M may contain the putative marmoset SEF. Further investigation employing ICMS and single unit electrophysiology in marmoset dorsomedial frontal cortex is required to fully investigate this putative homology.

We were also able to elicit saccades at rostral sites in area 46 and in anterior area 8aD. At these sites, saccades were evoked at high currents and long latencies, and did not exhibit any clear organization of direction or amplitude. As with our observations in other areas of marmoset frontal cortex, this finding is consonant with previous work in macaque (Robinson and Fuchs, 1969). Further investigation in the frontal pole of the marmoset brain is required to characterize this region.

Altogether, our data demonstrate a similar functional organization of the FEF in marmosets and macaques and provide a combined physiological characterization and anatomical localization that opens avenues for future exploration of FEF microcircuitry in marmosets. Electrophysiological studies in marmosets have the potential to complement ongoing work in the macaque model and human participants by advancing our understanding of laminar processes and their contributions to the oculomotor and cognitive functions of this area.

## Acknowledgements

We thank C. Vander Tuin, N. Hague, W. Froese, and K. Faubert for expert technical and surgical assistance, and care of the marmosets. This research was supported by the Canadian Institutes of Health Research grant FRN148365 to S.E. and the Canada First Research Excellence Fund to BrainsCAN.

## References

Avants BB, Tustison NJ, Song G, Cook PA, Klein A, Gee JC (2011) A Reproducible Evaluation of ANTs Similarity Metric Performance in Brain Image Registration. NeuroImage 54:2033–2044.

Awh E, Armstrong KM, Moore T (2006) Visual and oculomotor selection: links, causes and implications for spatial attention. Trends Cogn Sci 10:124–130.

Blum B, Kulikowski JJ, Carden D, Harwood D (1982) Eye Movements Induced by Electrical Stimulation of the Frontal Eye Fields of Marmosets and Squirrel Monkeys. Brain Behav Evol 21:34–41.

Bruce CJ, Goldberg ME, Bushnell MC, Stanton GB (1985) Primate frontal eye fields. II. Physiological and anatomical correlates of electrically evoked eye movements. J Neurophysiol 54:714–734.

Burish MJ, Stepniewska I, Kaas JH (2008) Microstimulation and architectonics of frontoparietal cortex in common marmosets (Callithrix jacchus). J Comp Neurol 507:1151–1168.

Corneil BD, Elsley JK, Nagy B, Cushing SL (2010) Motor output evoked by subsaccadic stimulation of primate frontal eye fields. Proc Natl Acad Sci U S A 107:6070–6075.

Elsley JK, Nagy B, Cushing SL, Corneil BD (2007) Widespread Presaccadic Recruitment of Neck Muscles by Stimulation of the Primate Frontal Eye Fields. J Neurophysiol 98:1333–1354.

Ferrier D (1875) The Croonian Lecture: Experiments on the Brain of Monkeys (second series). Philosopical Trans R Soc Lond 165:433–488.

Foerster O (1926) Zur operativen Behandlung der Epilepsie. Dtsch Z Für Nervenheilkd 89:137–147.

Ghahremani M, Hutchison RM, Menon RS, Everling S (2017) Frontoparietal Functional Connectivity in the Common Marmoset. Cereb Cortex 27:3890–3905.

Gilbert KM, Schaeffer DJ, Gati JS, Klassen ML, Everling S, Menon RS (2019) Open-source hardware designs for MRI of mice, rats, and marmosets: Integrated animal holders and radiofrequency coils. J Neurosci Methods 312:65–72.

Gottlieb JP, Bruce CJ, MacAvoy MG (1993) Smooth eye movements elicited by microstimulation in the primate frontal eye field. J Neurophysiol 69:786–799.

Heiney SA, Blazquez PM (2011) Behavioral responses of trained squirrel and rhesus monkeys during oculomotor tasks. Exp Brain Res 212:409–416.

Hung C-C, Yen CC, Ciuchta JL, Papoti D, Bock NA, Leopold DA, Silva AC (2015) Functional MRI of visual responses in the awake, behaving marmoset. NeuroImage 120:1–11.

Johnston KD, Barker K, Schaeffer L, Cutter DJ, Everling S (2018) Methods for chair restraint and training of the common marmoset on oculomotor tasks. J Neurophysiol 119:1636–1646.

Johnston KD, Ma L, Schaeffer L, Everling S (2019) Alpha Oscillations Modulate Preparatory Activity in Marmoset Area 8Ad. J Neurosci 39:1855–1866.

Knight TA, Fuchs AF (2007) Contribution of the Frontal Eye Field to Gaze Shifts in the Head-Unrestrained Monkey: Effects of Microstimulation. J Neurophysiol 97:618–634.

Li X, Morgan PS, Ashburner J, Smith J, Rorden C (2016) The first step for neuroimaging data analysis: DICOM to NIfTI conversion.

Liu C, Ye FQ, Yen CC-C, Newman JD, Glen D, Leopold DA, Silva AC (2018) A digital 3D atlas of the marmoset brain based on multi-modal MRI. NeuroImage 169:106–116.

Mitchell JF, Reynolds JH, Miller CT (2014) Active Vision in Marmosets: A Model System for Visual Neuroscience. J Neurosci 34:1183–1194.

Mott FW, Schuster E, Halliburton WD (1910) Cortical Lamination and Localisation in the Brain of the Marmoset. Proc R Soc Lond Ser B Contain Pap Biol Character 82:124–134.

Peterson J, Chaddock R, Dalrymple B, Van Sas F, Gilbert KM, Klassen LM, Gati JS, Handler WB, Chronik BA (2018) Development of a gradient and shim insert system for marmoset imaging at 9.4 T. In: Proceedings of the 26th Annual Meeting ISMRM, pp 4421. Paris, France.

Robinson DA, Fuchs AF (1969) Eye movements evoked by stimulation of frontal eye fields. J Neurophysiol 32:637–648.

Russo GS, Bruce CJ (1993) Effect of eye position within the orbit on electrically elicited saccadic eye movements: a comparison of the macaque monkey’s frontal and supplementary eye fields. J Neurophysiol 69:800–818.

Schall JD (1997) Visuomotor areas of the frontal lobe. In: Extrastriate cortex in primates, pp 527–638. Boston, MA: Springer. Available at: http://www.psy.vanderbilt.edu/faculty/schall/pdfs/VisuomotorAreasOfTheFrontalLobe-Ch13.pdf.

Schlag J, Schlag-Rey M (1987) Evidence for a supplementary eye field. J Neurophysiol 57:179–200.

Smith SM, Jenkinson M, Woolrich MW, Beckmann CF, Behrens TEJ, Johansen-Berg H, Bannister PR, De Luca M, Drobnjak I, Flitney DE, Niazy RK, Saunders J, Vickers J, Zhang Y, De Stefano N, Brady JM, Matthews PM (2004) Advances in functional and structural MR image analysis and implementation as FSL. NeuroImage 23:S208–S219.

Stanton GB, Deng S-Y, Goldberg EM, McMullen NT (1989) Cytoarchitectural characteristic of the frontal eye fields in macaque monkeys. J Comp Neurol 282:415–427.

Tomlinson RD, Bahra PS (1986) Combined eye-head gaze shifts in the primate. I. Metrics. J Neurophysiol 56:1542–1557.

Wakabayashi M, Koketsu D, Kondo H, Sato S, Ohara K, Polyakova Z, Chiken S, Hatanaka N, Nambu A (2018) Development of stereotaxic recording system for awake marmosets (Callithrix jacchus). Neurosci Res 135:37–45.

